# A Spatial and Temporal Transcriptomic Atlas of Mouse Intestinal Regeneration

**DOI:** 10.1101/2025.08.06.668727

**Authors:** Rachel Ofer, Tyler Tran, Ping He, Xia Qiu, Xinyu Hu, Emmanuel Zachariah, Curtis Krier, Ankit Saxena, Jiekun Yang, Michael P. Verzi

## Abstract

**Background & Aims:** The intestinal epithelium exhibits a remarkable capacity for regeneration following injury. However, the spatial and temporal dynamics of the injury-repair cycle remain incompletely understood.

**Methods:** We employ spatial transcriptomics to create an atlas of the damage and repair response to ionizing radiation in the mouse intestine. We map molecular events driving epithelial recovery over a six-day period and 23 biological samples, spanning the early apoptotic response to tissue remodeling and repair.

**Results:** The datasets capture mRNA of 19,042 genes in ∼26 million bins at 2µm resolution. Analysis revealed transcriptional patterns and niche signals that would remain undetected in bulk or single-cell approaches, including a non-random activation of interferon-target genes. Temporal shifts in cytokine and growth factor gene expression, particularly in the crypt and lower villus regions, corroborate published studies and reveal new predictions of the mechanisms governing intestinal healing. Global transcriptional upregulation was observed in the regenerating epithelium, suggesting hypertranscription is a hallmark of intestinal repair. Furthermore, we observe altered cellular differentiation trajectories and villus patterning at the early stages of regeneration.

**Conclusions:** Together, our work provides a detailed spatiotemporal map of intestinal regeneration at subcellular resolution and nearly whole-genome scale. These data lay the groundwork for future discoveries and therapeutic strategies to enhance epithelial repair in inflammatory bowel diseases and other gastrointestinal pathologies or in response to side-effects of cancer therapies.

## INTRODUCTION

The intestine provides one of nature’s best examples of how the evolution of structure is associated with function. The elegant spatial composition of the repeating pattern of crypts and villi provides immense surface area to facilitate absorption of nutrients and water, while the vast surface area also in turn presents a large interface with microbial communities and is accompanied by an active mucosal immune system. The highly proliferative nature of the tissue also makes it susceptible to damage. During acute intestinal injury from pathogens, chemotherapies, radiation therapies, or inflammatory diseases, intestinal homeostasis is disrupted, triggering a cascade of regenerative responses. Damaged epithelial cells initially undergo repair or cell death, followed by immune cells and stromal fibroblast providing regenerative signals, and finally regenerative stem cells undergo hyperproliferation to recover lost epithelium. Ultimately, the damaged tissue is remodeled to restore the spatial organization of the homeostatic intestine^1–4^.

New technologies enabling in situ profiling of gene expression^5, 6^ are beginning to shed light on the damage/repair cycle in the intestine. In models of mouse colitis, spatially resolved transcriptomic studies have profiled subsets of genes to make discoveries such as alterations in stromal fibroblasts^7^ or spatially restricted signaling shifts in the regenerating epithelium^8^. Gene panels have been probed in situ to uncover the role of CD8 T cells during viral infection of the small intestine^9^. Radiation therapy is a pillar of cancer treatment but causes enteropathy and reduced quality of life^10^. In mice, ionizing radiation (IR) causes acute epithelial damage and crypt loss, triggering surviving epithelial cells to enter a regenerative state^11^. Regenerating cells shift their transcriptional programs to meet the demands of rapid expansion to restore homeostasis^1, 12, 13^. IR-induced injury in mice is a valuable model for studying epithelial regeneration^14^. However, substantial knowledge gaps remain regarding the spatial and temporal dynamics in the transcriptomes of cells that initiate and drive this regenerative response.

Recent technological advances allow nearly the whole transcriptome to be detected at a resolution of 2μm bins^15^. We generated four high-resolution spatial transcriptomics datasets across 23 biological replicates spanning 0-6 days post-IR. This approach enabled simultaneous profiling of almost the full genome at near single-cell resolution. By capturing spatial and temporal gene expression dynamics, the data reveals changes in cellular behavior, tissue architecture, and cell-cell interactions during intestinal regeneration, including shifts in the mechanics of epithelial differentiation, activation of hypertranscription by regenerating epithelium, and an altered stromal signaling environment during repair. The reproducibility across independent experiments and easy data accessibility provide a robust and valuable resource (<u>GSE303705</u>) for studying epithelial repair and tissue remodeling.

## MATERIALS AND METHODS

### Animals

Mouse protocols and experiments adhered to ARRIVE standards, were approved by the Rutgers Institutional Animal Care and Use Committee and all relevant ethical regulations were followed, including efforts to reduce the total animals necessary. C57BL/6J (JAX #000664) male littermates (8-12 weeks old) were used for spatial transcriptomics after whole-body IR at 12 Gy using a cesium-137 irradiator. Tissues were collected between 10am-2pm.

### Tissue processing for Visium HD Spatial Transcriptomics

Jejunal tissues were washed with cold PBS and opened longitudinally. Tissues from different treatments were arranged together for co-embedding and fixed overnight with 4% paraformaldehyde at 4°C. Tissues were dehydrated in ascending alcohols, processed with xylene, and embedded, generating a total of 4 formalin-fixed paraffin-embedded (FFPE) tissue blocks and cut at 5-μm thickness. H&E stain and imaging were performed according to Visium HD FFPE Tissue Preparation Handbook (CG000684) and guides (CG000518 revision C and CG000688). All tissue samples passed a DV200 quality check for RNA integrity. Imaging was done in Olympus VS120 format.

### Visium HD Spatial transcriptomics sequencing

Samples were processed using the Visium CytAssist platform and sequenced following the Visium HD Spatial Gene Expression Reagents Kits User Guide (CG000685). Libraries were prepared using the 10x Genomics Visium HD FFPE protocol and sequencing was performed on an Illumina NovaSeq X Plus platform using paired-end 150-bp reads with dual indexing (10-bp index length). Libraries were sequenced on partial lanes with a 5% PhiX control spike-in for run quality calibration. Sequencing data were demultiplexed by the provider. Total sequencing output per replicate can be found in Supplemental Fig. 1B.

### Visium HD Spatial Gene Expression Analysis

Space Ranger (v3.1.3 and v4.0) was used to process Visium HD libraries, outputting data binned at 2µm and 8µm resolution. Unless otherwise noted, downstream analyses were conducted using the 8µm resolution data.

### Clustering and differential gene expression (DGE)

Graph-based and k-means clustering were performed within Loupe Browser (v8.0). In some cases, sub-clustering of specific tissue regions was performed with the “reanalyze” feature. Unless specified, DAVID Gene Ontology (GO) terms^16, 17^ were based upon differentially expressed genes filtered for Log_2_Fold-Change > 1, mean normalized average (MNA) > 0.03, and adjusted *P val* < 0.05, performed within Loupe Browser (v8.0). For epithelial-specific analysis (Fig. 2) unsupervised k-means clustering identified five distinct clusters. Stromal bins, identified through graph-based clustering, were excluded prior to downstream analysis. DGE was conducted within the day 0 dataset to compare the five epithelial clusters, using the same gene filtering criteria. Morpheus (https://software.broadinstitute.org/morpheus/) was used to visualize normalized transcript levels of genes of interest. For stromal specific analysis (Fig. 4), stromal clusters were first identified using graph-based clustering. To resolve stromal heterogeneity, the “reanalyze” feature was used to define three sub-clusters: villus stroma, inter-stroma, and sub-stroma. Equivalent subclusters were identified across timepoints, and DGE was performed. To focus on stromal expression, epithelial-associated genes (n = ∼10,000) were derived from WT epithelium RNAseq, with genes having FPKM>1 in at least 2 of the 3 samples (GSE112946)^18^ removed before analysis.

**Figure 1.**
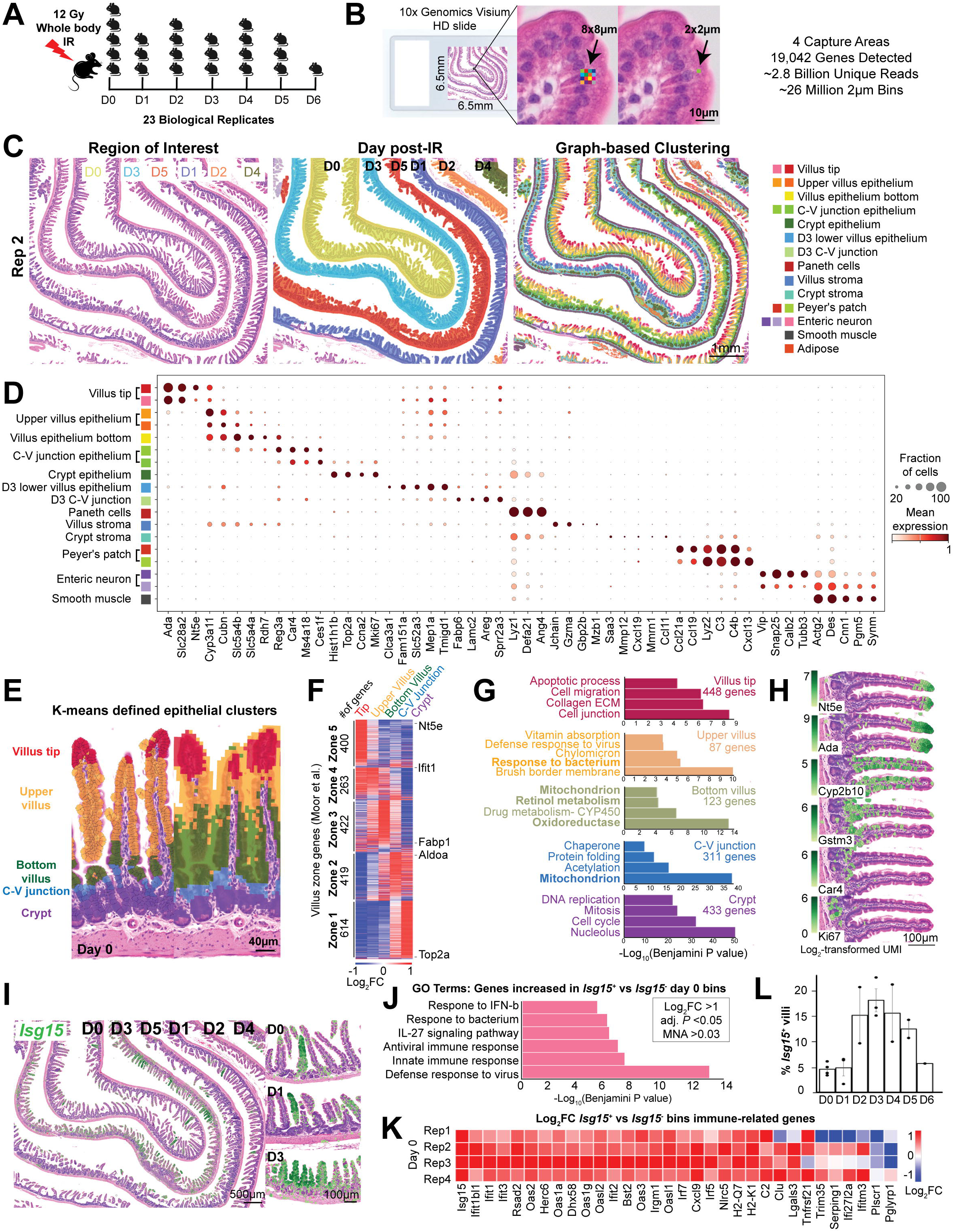
A spatial transcriptomic atlas of post-irradiation intestinal regeneration. (**A**) Schematic of VisiumHD spatial transcriptomics (ST) experimental design. Regions of interest (ROI) = 4, totaling 23 mice (n = 1-5 per timepoint). (**B**) High-resolution spatial transcriptomics generated data at 2×2 µm bin resolution. For some downstream analyses, data were aggregated into 8×8 µm bins. Across all replicates, ∼2.8 billion unique reads and ∼26 million 2µm bins were analyzed, detecting 19,042 genes. (**C**) Representative spatial capture region from replicate 2; highlighted by timepoints (middle panel) and with unsupervised graph-based clustering of 8µm bins (right panel), revealing distinct spatial regions along the crypt-villus axis and dynamic transcriptomes during intestinal regeneration. (**D**) Mean normalized log_2_-transformed expression values for top upregulated genes for the spatial graph-based clusters. (**E**) High-resolution spatial clustering identifies five transcriptionally distinct epithelial clusters along the crypt-villus axis in non-irradiated tissue, visualized with cell segmentation (left) or 8µm bins (right). Bins associated with stromal clusters were removed. (**F**) Log_2_FC expression of published zone-specific gene sets^25^ across the five epithelial clusters aligns with the current data set (Fig. 1E), validating the functional identity of each cluster. (**G**) GO enrichment analysis of epithelial clusters at day 0 reveals zone-specific biological processes along the crypt-villus axis. DGE was performed between the five epithelial clusters to generate gene lists for each zone (Log_2_FC>1, *P* < 0.0.5, Average >0.03). Enriched GO terms reflect known functional specialization. GO enrichment was calculated using DAVID^16, 17^, with significance defined by adjusted *P* < 0.05 (Benjamini-Hochberg correction). (**H**) Expression of representative markers visualized in Loupe Browser confirms spatial enrichment patterns. (**I**) Spatial transcriptomic mapping of interferon signaling genes reveals region-specific expression patterns during homeostasis and regeneration. Log_2_-transformed expression of *Isg15*, with cell segmentation highlighting *Isg15*^+^ villi of day 0,1 and 3 post-IR. (**J**) GO enrichment analysis for *Isg15*^+^ vs *Isg15*^-^ bins on day 0 indicates upregulation of interferon-associated pathways. (**K**) DGE of interferon-stimulated genes (ISGs), shown as log_2_-fold-change between *Isg15*^+^ and *Isg15*^-^ bins in day 0 tissues across all replicates. Analysis done using 8µm bins. (**L**) Percentage of villus-associated bins expressing *Isg15* across all replicates from day 0 to day 6. n = 4(D0), 3(D1), 2(D2), 3(D3), 2(D4), 2(D5), 1(D6), n=17 ** methods for figures J and K: Smooth muscle, enteric neuron, crypt and crypt-stroma bins were excluded from analysis. DGE was run between *Isg15*^+^ and *Isg15*^-^ bins. *Isg15* gene set was inserted and Log_2_FC values displayed in the heatmap. Expression profile of immune-related genes populated from RNA-seq and GO analysis, from five GO terms: GO:0051607, GO:0009615, GO:0002376, GO:0045087, and GO:0045071.^27^

**Figure 2.**
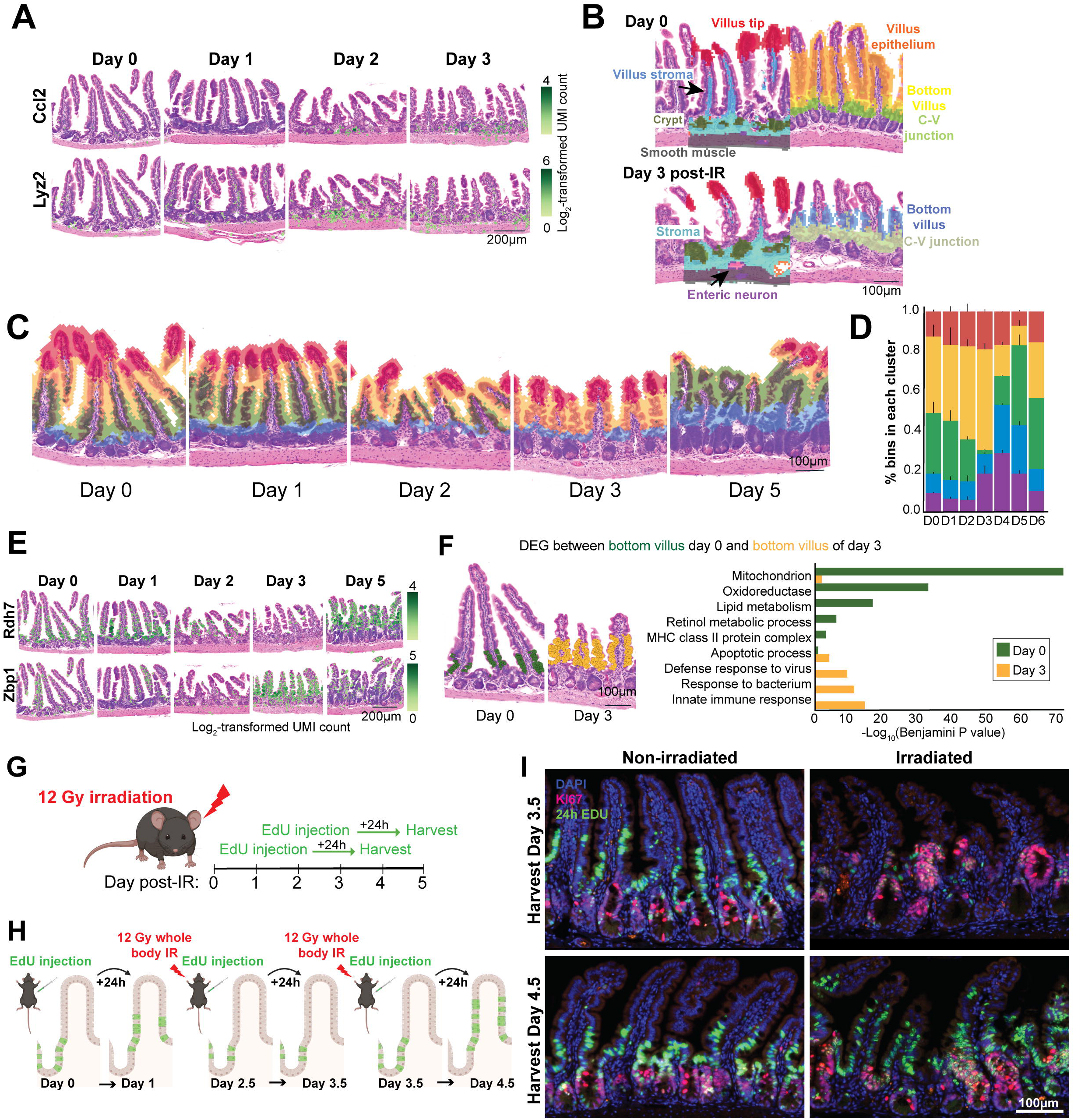
The villus bottom epithelium exhibits a transcriptional shift towards maturity and innate immunity at day 3 post-IR. **(A)** Increased recruitment of monocytes from day 0 to day 3 post-IR, visualized via Log_2_-transformed expression of macrophage markers *Ccl2* and *Lyz2*. **(B)** Unsupervised clustering identifies similar (villus tip, enteric neurons, crypt, and smooth muscle) and distinct (bottom villus epithelium, C-V junction) transcriptional profiles for the same regions on day 0 and day 3 post-IR. **(C)** The cluster representing early post-mitotic villus cells (green on day 0) is specifically reduced or absent at day 3 post-IR, instead replaced by a cluster associated with upper villus at day 0 (yellow). Cluster assignments were derived from k-means clustering restricted to epithelial bins. **(D)** Percent distribution of epithelial clusters at day 0 and day 3 post-IR. Values reflect the proportion of epithelial bins assigned to each cluster normalized to the total number of epithelial bins per sample. n = 5(D0), 3(D1), 3(D2), 3(D3), 2(D4), 3(D5), 1(D6). Error bars represent SEM across biological replicates. **(E)** Spatial-temporal expression (log_2_-transformed UMI counts) of representative markers (*Rdh7, Zbp1*) that exhibit a shift in the post-IR timepoints. **(F)** DGE between bottom villus cluster of day 0 and bottom villus cluster of day 3. Representative GO Terms. **(G)** Schematic of EdU-based epithelial proliferation assay. Mice were administered EdU (5-ethynyl-2′-deoxyuridine) at 2.5- and 3.5 days post-IR, and tissues were collected 24 hours later to track the proliferation and migration of newly replicating cells. **(H-I)** EdU staining reveals retention of proliferative epithelial cells in lower villus and crypt regions between days 2.5-3.5 post-IR, followed by resumed upward migration by day 4.5 post-IR. This pattern suggests a delay in contribution of new cells to the villus base during regeneration. Nuclei were counterstained with Hoechst 33342. n=2.

**Figure 3.**
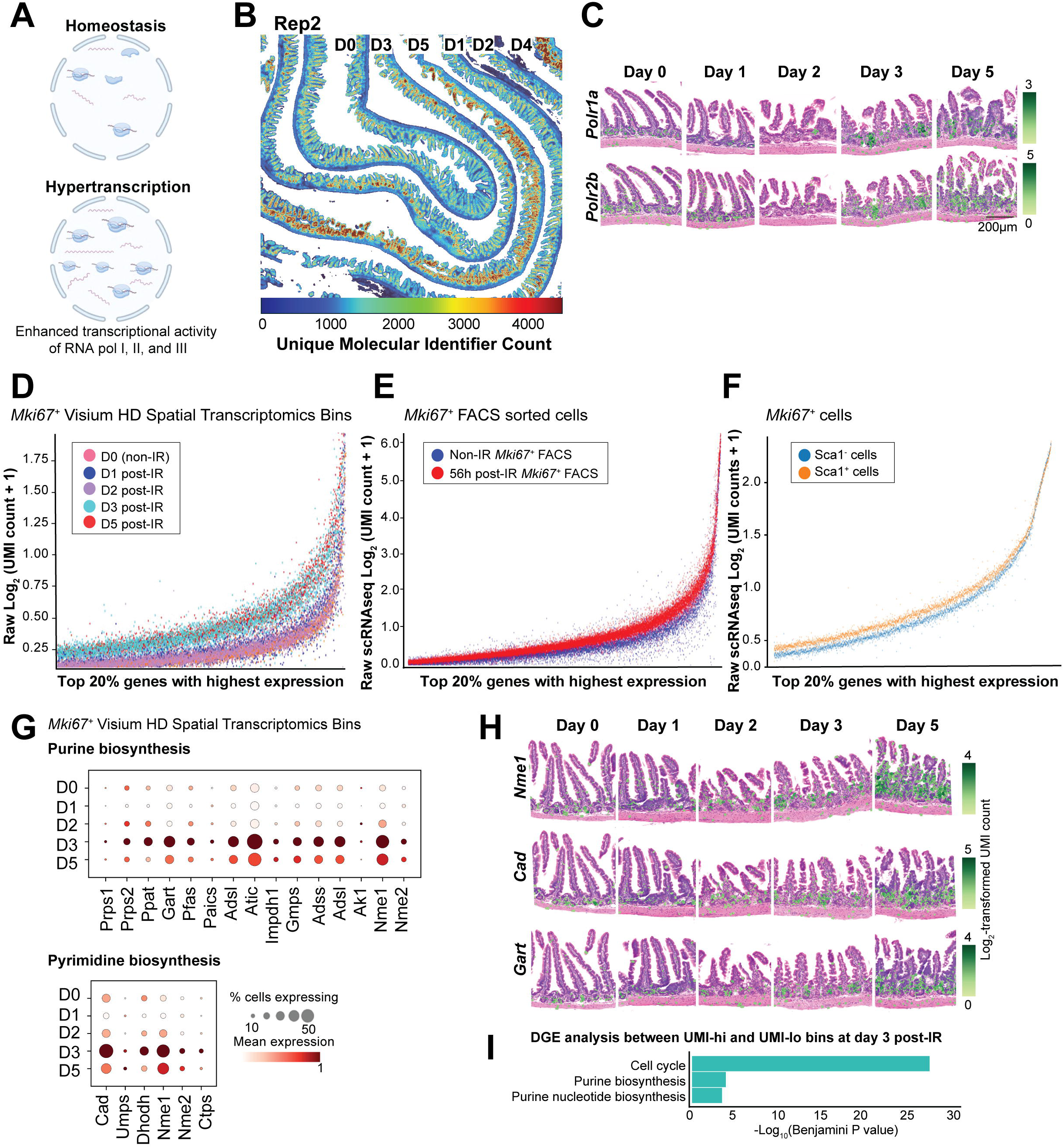
Global transcript levels are elevated in regenerating intestinal tissues, indicating hypertranscription is a feature of intestinal regeneration. **(A)** Enhanced activity of RNA polymerases I, II, and III marks hypertranscription in regenerating intestinal tissues. **(B)** Heatmap of raw unique molecular identifier (UMI) counts demonstrates increased transcript abundance at days 3 and 5 post-IR, with elevated total UMI counts per barcode (replicate 2). **(C)** Spatial expression of RNA polymerase subunits shows upregulation of *Polr1a* and *Polr2b* across regenerative tissues. Log_2_-transformed expression, replicate 2. **(D–F)** Regenerating intestinal tissues show global transcriptomic amplification in proliferative epithelial cells. Plots depict mean log_2_-transformed raw unique read counts of the top 20% most highly expressed genes across post-IR timepoints in: (D) Spatial transcriptomic data (Replicate 2), filtered for proliferative epithelial bins (*Mki67*^+^ and either *Epcam^+^* or *Vil1*^+^); (E) FACS-sorted *Ki67-RFP*^+^ cells at 56 hours post-IR compared to non-IR *Ki67-RFP*^+^ cells from Chen et al. (GSE222505)^20^; (F) *Mki67*+ cells subset from sorted *Sca-1^+^* and *Sca-1^−^* epithelial populations post-helminth infection, from Nusse et al. (GSE108233)^21^. **(G)** Nucleotide metabolism genes are upregulated in proliferative epithelial bins (*Mki67^+^*) from (replicate 2) beginning at day 3 post-IR. Normalized mean expression of key genes demonstrates increased *de novo* purine and pyrimidine biosynthesis to support regenerative demands. **(H)** Elevated expression of nucleotide synthesis genes (*Cad, Nme1, Gart*), localized to proliferative cell compartments. Visualized with cell segmentation (replicate 2). **(I)** GO enrichment analysis of genes upregulated in transcriptionally active regions (UMI-high bins) at day 3 post-IR reveals enrichment of biosynthetic and RNA processing pathways. Bins with >2500 UMI counts were classified as “UMI-high,” and <2500 as “UMI-low.” Differential expression was performed between high and low bins within each timepoint (criteria: log_2_ fold-change > 1, MNA > 0.03, adjusted *P* < 0.05). Day 3 post-IR: 177 genes in UMI-high vs 427 in UMI-low bins were analyzed.

**Figure 4.**
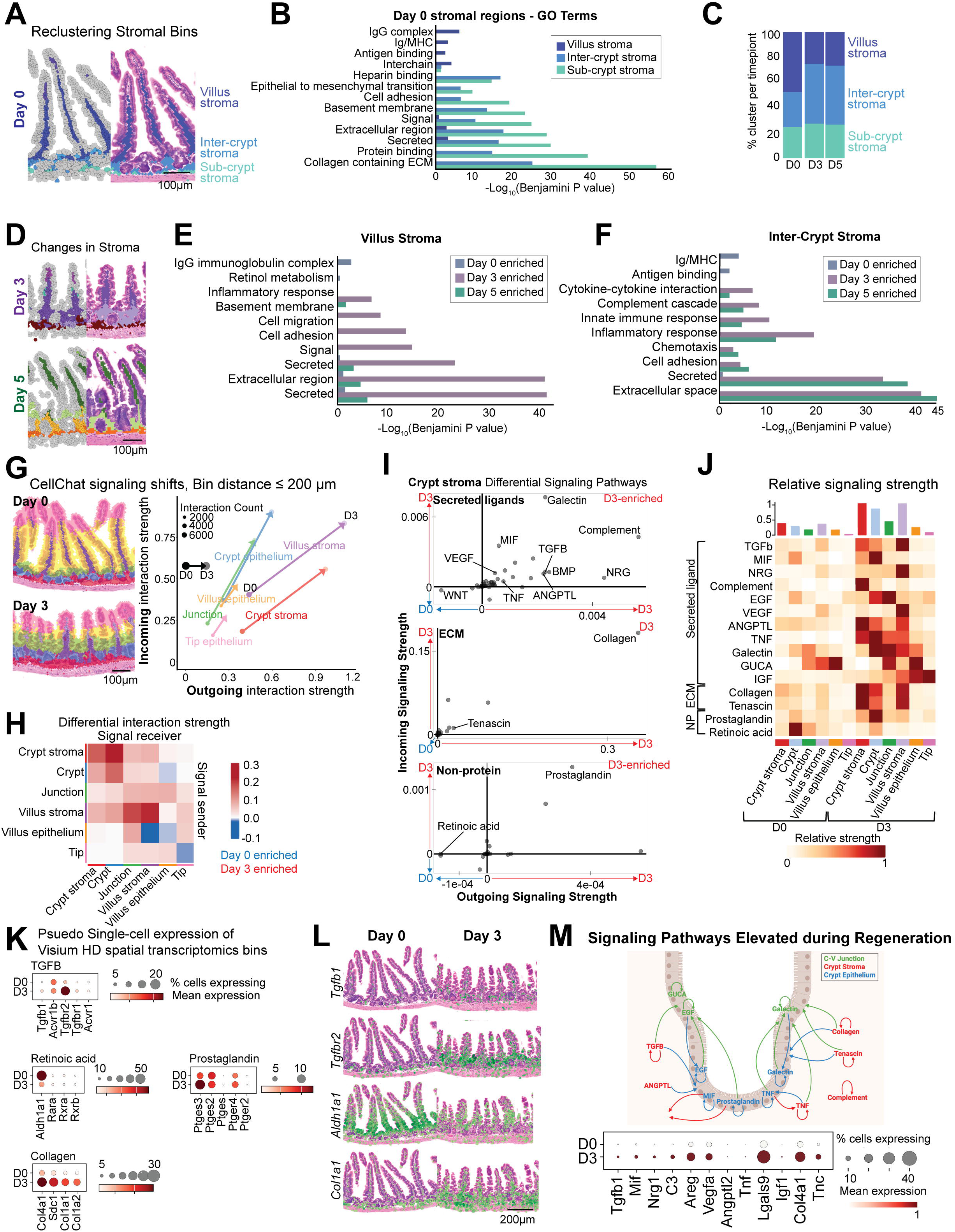
Spatial transcriptomics reveals regional signaling dynamics during regeneration and provides insight into stromal-epithelial interactions. **(A)** Stromal regions were defined using unsupervised graph-based clustering of spatial transcriptomics data (replicate 2 shown). Clusters include villus stroma (dark blue), inter-crypt stroma (light blue), and sub-crypt stroma (turquoise), used for subsequent analyses in panels B-F. **(B)** GO enrichment analysis in non-irradiated (day 0) control tissue reveals distinct biological functions for each stromal domain, including structural support, immune surveillance, and signaling regulation. **(C)** Proportional representation of each stromal region (based on number of bins in each cluster) across days 0, 3, and 5 post-IR, reflecting dynamic tissue remodeling. **(D)** Stromal domains identified at day 3 and day 5 post-IR using graph-based clustering. Villus stroma is shown in dark purple (day 3) and dark green (day 5); inter-crypt stroma in light purple (day 3) and light green/orange (day 5). These domains were used for GO term analyses in panels E and F. **(E)** GO term enrichment in villus stroma shows upregulation of immune activation and tissue repair pathways at days 3 and 5 post-IR relative to control. **(F)** Inter-crypt stroma shows a shift in enriched pathways toward innate immune responses and tissue remodeling during regeneration. **(G)** Spatial CellChat analysis comparing day 0 and day 3 identifies enhanced overall signaling activity post-IR. Left: Cell domains defined by unsupervised clustering. Right: Changes in outgoing vs. incoming signaling strength for each region, based on CellChat predictions (replicate 2). **(H)** Spatial map of differential signaling strength shows enhanced signaling in stromal regions at day 3 post-IR (red) compared to day 0 (blue), measured by incoming and outgoing activity. **(I)** Differential signaling activity in the crypt stroma between day 0 and day 3 post-IR, assessed using CellChatDB’s secreted ligand, non-protein, and ECM-receptor interaction databases. Highlighted pathways include those previously implicated in epithelial repair and stromal remodeling. Pathways enriched at D3 are in the upper right quadrant. **(J)** Quantification of total signaling strength (incoming + outgoing) across stromal regions for selected pathways known to be involved in intestinal regeneration. **(K)** Dot plots of normalized log_2_-transformed expression levels of key ligands and receptors in TGFB, collagen, retinoic acid and prostaglandin signaling pathways across regions defined in panel G. **(L)** Log_2_ expression of selected TGFB, collagen, and retinoic acid signaling genes in day 0 and day 3 tissues demonstrates shifts in pathway activation during regeneration. **(M)** Spatially restricted CellChat predictions recapitulate known regeneration-associated signaling pathways and are supported by corresponding changes in ligand and receptor gene expression. Pseudo-single cell analysis for these pathways is shown via dot blot below.

### Cell compartmentalization

Nuclear and cellular segmentation was conducted using Space Ranger (v4.0) and visualized on Loupe Browser (v9.0). This method partitions barcodes into bins based on automated nuclei detection in the corresponding image. The method matched H&E-stained microscopy images to accurately associate spatial transcriptomic data.

### EdU pulse-chase

Male and female littermate mice were injected with 200µg EdU by intraperitoneal injection. Tissues were processed, embedded in paraffin, and sectioned as described above. Antigen retrieval for KI67 was performed using pH 6 citrate-based heat-induced epitope retrieval. Tissues were blocked using 3% BSA in PBS. EdU was detected by click chemistry reaction according to manufacturer’s instructions from the Invitrogen™ Click-iT™ EdU Cell Proliferation Kit for Imaging, Alexa Fluor™ 488 dye (C10337). Tissues were incubated with primary antibody overnight at 4°C: Ki67 (Abcam ab16667, 1:400 dilution in 3% BSA in PBS) then Goat anti-Rabbit IgG (H+L) Cross-Adsorbed Secondary Antibody, Alexa Fluor™ 555 (Invitrogen, 1:1000 in 3% BSA) was used for visualization. After washing, nuclei were counter-stained with DAPI (Biotium 40043). Stained tissues were prepared in mounting media and imaged using Leica THUNDER microscope and Leica LASX acquisition software with the same parameters across all tissues. ImageJ and LASX were used to adjust contrast and brightness. When adjustments of sharpness, contrast, or brightness were made, they were applied uniformly across comparative images.

### Raw counts scatterplot analysis

To assess hypertranscription, 8µm Space Ranger output bins were filtered in Scanpy^19^ for expression of *Mki67* and either *Vil1* or *Epcam*, isolating proliferative epithelial cells only. Average gene raw counts were calculated, log_2_-transformed, and sorted by mean expression across samples. To compare mRNA abundance between regenerative and homeostatic states, scRNA-seq datasets from Chen et al.^20^ (GSE222505; Ki67+ FACS sorted, 56h post-IR) and Nusse et al.^21^ (GSE108233; Sca-1^+^ and Sca-1^-^ crypt cells from *H.polygyrus* infected mice) were analyzed. Sca-1 dataset was further filtered for *Mki67^+^* expression. Scatterplots were created as described for the spatial data. Dot-plots of gene-specific expression were created using Scanpy^19^ with normalized counts. Spatial bins required a minimum of 40 transcripts and 10 genes; scRNA-seq cells required 1,000 transcripts and 100 genes.

### Cell-cell signaling analysis

CellChat v2 (2.1.2) was used to analyze bin-to-bin communications^22^ from 8µm Space Ranger output. Bins with ≥40 unique molecular identifiers (UMIs) were selected based on treatment timepoint and cell type region, as identified by graph-based clustering, and exported to Seurat. Normalized counts and spatial coordinates were selected for input into a CellChat object^23^. Analyses used the Secreted Signaling, ECM-Receptor, Cell-Cell Contact, and Non-Protein databases from CellChatDB. The function “computeCommunProb” was used to predict communications and their strengths between the regions, with a truncated mean value set to trim 0.001, interaction range limited to 200µm, and contact range limited to 20µm. A minimum threshold of 10 cells in each group was required to detect signaling, and “computeCommunProbPathway” was used to identify the signaling pathways involved. For differential comparisons, the workflows for Day 0 and Day 3 clusters were analyzed separately, then merged to compare differentially enriched signaling pathway communication strengths. Relative communication strengths were visualized with ComplexHeatmap^24^.

### Quantification and statistical analysis

p-values are adjusted using the Benjamini-Hochberg correction for multiple tests and generated using Loupe Browser (v8.0 and v9.0). Fold-change was calculated as the ratio of normalized mean gene UMI counts in each cluster relative to all others. For supplemental Fig. 1J, villi of each timepoint and replicate were marked as *Isg15^+^* or *Isg15^−^* and converted to binary strings, with villus units representing distance. The normalized Ripley L-function and 95% confidence intervals from 1,000 Monte Carlo simulations were calculated and graphed.

## RESULTS

### A Single-Cell Spatial Transcriptomic Resource for Intestinal Regeneration

Intestinal damage from ionizing radiation (IR) is a limiting factor for radiation therapy and a widely used model to understand the intestinal damage-repair cycle. To better appreciate the spatial and temporal response of the intestine to ionizing radiation, we conducted a spatial transcriptomics time-series of mouse small intestine after exposure to 12Gy IR. Jejunal segments from 23 independent biological replicates spanning days 0 through 6 post-IR (Fig. 1A) were embedded into 4 paraffin blocks and processed using the VisiumHD platform (Fig. 1B-C, Supplemental Fig. 1A). VisiumHD uses a continuous array of 2um barcoded probe-capture grids to determine RNA expression in the tissue with high spatial resolution and near-genome scale breadth^15^. A representative bin is shown to illustrate sub-nuclear resolution (Fig. 1B). Across the 4 capture areas, 19,042 genes were detected spanning ∼26 million 2um bins at a sequencing depth of ∼2.8 billion unique reads (Fig. 1B, Supplemental Fig. 1A-B). Unsupervised clustering of the bins revealed striking patterns of gene expression, consistent with known spatial patterns in intestinal biology (Fig. 1C-D). For example, zonation of the villus epithelium^25^ was apparent, with villus tip, middle, and bottom regions clustering separately (Fig. 1E). Distinct bins expressing markers of smooth muscle or enteric glial cells (*Gfap*) comprised clusters localized at the periphery (Fig. 1D and Supplemental Fig. 1C). Stromal cell populations were detected in the villus core and inter-crypt regions and expressed genes associated with immune cell populations and fibroblasts (Fig. 1D, Supplemental Fig. 1D-E). The data was probed to detect canonical marker genes; for example, markers of B cells (*Jchain*), T cells (*Cd8a*), M cells (*Gp2*), Tuft cells (*Avil*), proliferative cells (*Mki67*), and regenerative stem cells (*Ly6d, Anxa1, Anxa8, Sprr1a, Tead4, Tnfrsf12a, and Clu)* were visualized in space and time (Supplemental Fig. 1D-G).

Unsupervised k-means clustering of bins’ transcriptomes spanning the crypt-villus axis identified five distinct epithelial zones: villus tip, upper villus, bottom villus, crypt-villus junction, and crypt epithelium (Fig. 1E). The distinct domain of the crypt-villus junction was reliably preserved in the current study unlike in other studies using conventional cell isolation methods, where most epithelial preparations are damaged at the crypt-villus junction, potentially compromising the results. DGE analysis (Log_2_FC>1,adj. *P*<0.05, MNA>0.03) between these epithelial zones yielded gene sets characteristic of each region (Supplemental Table 1), and the transcriptomes of each zone correlated strongly with previously described villus zonation signatures^25, 26^ (Fig. 1F). Zonation patterns were consistent across independent experiments (Supplemental Fig. 1H). GO Term analysis corroborated established functions of epithelial zonation^25^ (Fig. 1G, Supplemental Table 1). Examples of spatially localized expression of zone-specific genes along the crypt-villus axis is shown in Fig. 1H.

Finally, unique features of intestinal biology that might be missed by single-cell approaches could be captured in the spatial datasets. For example, a baseline level of interferon signaling appeared in control animals in an intriguing spatial pattern. Roughly 5% of villi exhibited elevated marker genes of interferon pathway activation, such as *Isg15*. These *Isg15-*hi villi appeared sporadically distributed along the tissue of non-IR samples (Fig. 1I). To better appreciate this pattern, we compared expression of genes in *Isg15^+^* bins with *Isg15^−^* bins of the villus epithelium (Fig. 1J, Supplemental Table 2). This analysis revealed a whole battery of interferon-stimulated genes^27^ that are elevated in *Isg15*-hi villi (Fig. 1K). During the damage-repair cycle, increasing numbers of villi exhibited interferon-stimulated gene expression (Fig. 1L), and these villi tended to cluster in groups as supported by Ripley’s L function, significant in day 3 timepoints (Supplemental Fig. 1J), hinting at potential communication between adjacent villi to allow the signal to spread (Fig. 1I, Supplemental Fig. 1I-J). Co-expression analysis revealed that most bins expressing one IFN-pathway target also expressed others (Supplemental Fig. 1I). Thus, we provide a resource for the field to explore, generate, and test hypotheses concerning the mechanisms controlling intestinal damage and repair. We present these data as an easily downloadable and accessible dataset (<u>GSE303705</u>) and provide an accompanying Supplemental ‘spatial data guide’ for easy use.

### The villus bottom epithelium exhibits a transcriptional shift towards maturity and innate immunity at day 3 post-IR

Dynamic expression of cell lineage markers could be observed throughout the regenerative timeline. For example, macrophage recruitment to damaged crypts at days 2-3 post-IR can be visualized through the dynamic expression of monocyte recruitment genes (*Ccl2*) and macrophage marker (*Lyz2*) genes (Fig. 2A). While some clusters remained relatively stable throughout the intestinal damage/repair cycle, such as those corresponding to the villus tip, smooth muscle, and enteric nervous system (Fig. 2B, left half), other regions exhibited dynamic transcriptional profiles. Clusters associated with the crypt-villus junction and the bottom villus epithelium showed enough transcriptional change to be reassigned to new clusters during the regenerative phase (Fig. 2B, right half).

On day 3 post-IR, the transcriptional profile of villus bottom bins diverged markedly from other timepoints; instead of clustering with other villus bottom bins, they shifted to a cluster typically associated with the upper villus epithelium of days 0 and 5 (day 3 yellow cluster, Fig. 2C-D). This suggests that, despite being spatially located at the villus base, these day 3 cells exhibit a transcriptomic signature characteristic of more mature epithelial cells. This regenerative shift in epithelial zonation was reproducibly observed in independent replicate experiments (Supplemental Fig. 2A).

On day 3 post-IR, bins at the villus bottom expressed most “mature” yellow cluster-defining genes (Supplemental table 1), but also upregulated genes (Log_2_FC > 1, adj. *P*<0.05, MNA>0.03) associated with innate immune functions (*Zbp1* marker, Fig. 2E; Supplemental Fig. 2B, Supplemental table 3 GO Terms), a characteristic typically enriched to the upper villus epithelium under homeostatic conditions (Fig. 1G). These findings indicate that the villus bottom epithelium at day 3 exhibits transcriptional features of more mature upper-villus epithelium, along with heightened expression of immune-related genes, likely reflecting the inflammatory environment associated with tissue injury at this timepoint.

Consistent with these observations, DGE comparing day 0 villus bottom bins (green cluster, Fig. 2F) with those from day 3 (yellow cluster, Fig. 2F) showed a near-complete loss of genes that define the homeostatic bottom villus cluster at day 3 (Supplemental table 4). 76 genes normally enriched at the villus bottom were significantly downregulated in the day 3 villus bottom and were associated with functions such as prostaglandin and retinol metabolism, mitochondrion, and oxidoreductase (Supplemental Fig. 2C, Supplemental table 5). A representative example is *Rdh7*, a gene involved in retinol metabolism, which is markedly downregulated at the day 3 villus bottom (Fig. 2E), highlighting the dynamic events at this region.

A potential hypothesis for why cells at the villi base on day 3 post-IR were associated with a mature state is a change in the contribution of newly differentiated cells from the crypt to the villus post-IR. To explore whether migration of cells from the crypt onto the villus is altered post-IR, we conducted EdU pulse-chase experiments (Fig. 2G-H). Based upon previous work^28^, proliferation resumes around 2.5 days post-IR; we conducted 24-hour pulse-chase experiments at 2.5- and 3.5-days post-IR. EdU-labeled cells from mice injected at day 2.5 post-IR and harvested at day 3.5 post-IR remained in crypts (marked by *Ki67*), instead of contributing to the villi as occurs during homeostasis (Fig. 2I). This finding indicates a zone of proliferating cells that do not contribute to the villus base occurs around day 3, and the transcriptomes indicate the cells at the base of the villus continue to mature. In contrast, cells labeled with EdU injection at 3.5 days post-IR contribute progeny cells up onto the villi (Fig. 2I), resuming the pattern of homeostasis. We therefore conclude that there is a pause in contribution of new villus cells from the crypt at the day 3 timepoint, and this pause leads to acquisition of a more mature transcriptome within the villus bottom at day 3 post-IR. Similar shifts in the transcriptome at the villus bottom during regeneration were seen across replicates (Supplemental Fig. 2D-F, Supplemental tables 6,7 and 8).

### Spatial Time-course Provides Evidence for Hypertranscription in the Regenerating Epithelium

Hypertranscription is the phenomenon in which cells elevate their transcriptional output, often associated with stem cell activity^29–32^ (Fig. 3A). Normalization methods in techniques such as RNA-seq and scRNA-seq usually equilibrate total mRNA counts between samples or cells, masking the detection of hypertranscription^30^. As part of quality control during spatial data analysis, we examined whether unique molecular identifier (UMI) count distributions were uniform across the capture area. While the UMI distributions passed quality control for an even distribution, we noticed an interesting pattern of elevated UMI counts localized to the regenerating epithelium, particularly at days 3-5 post IR (Fig. 3B), an observation that was consistent across replicates (Supplemental Fig 3A).

A hallmark of hypertranscription is the elevated expression of RNA polymerases^33^, which we also observed in the regenerating epithelium (Fig. 3C). Other genes associated with hypertranscription, such as *Chd1* and *Myc*^34, 35^, were similarly enriched in the regenerating epithelium (Supplemental Fig. 3B). To determine whether the increase in transcript levels was due to the upregulation of specific genes or a genome-wide increase, we analyzed the spatial transcriptomics bins as pseudo-single-cell data. *Ki67*-expressing bins were extracted at each timepoint to allow direct comparison of proliferating cells across time points. Interestingly, raw

UMI counts were elevated across most expressed genes in regenerative timepoints compared to non-IR controls (Fig. 3D), consistent with hypertranscription. We sought additional evidence for hypertranscription using independent methodologies; scRNA-seq of cells sorted from Ki67-RFP knock-in mice^36^ of non-IR controls compared to IR tissues^28^ (Fig. 3E), and proliferating cells (*Mki67^+^)* that are either expressing the regenerative marker Sca1 or not (*Ki67*^+^Sca1^+/-^)^21^ (Fig. 3F). Both analyses showed a similar pattern: UMI count was elevated across most genes in the regenerating tissues (Fig. 3E-F) compared to proliferating cells from homeostatic tissues.

Hypertranscription necessitates increased nucleic acid production. Correspondingly, transcript levels for genes involved in *de novo* nucleotide metabolism, including both purine and pyrimidine synthesis pathways^37^, were upregulated in the proliferating regions of the regenerating tissues compared to non-IR controls (Fig. 3G-I, Supplemental table 9). These observations were consistent across independent experiments (Supplemental Fig. 3C-D). Thus, as observed in various stem and progenitor populations^30^, spatial transcriptomics data revealed that the regenerating epithelium enters a state of hypertranscription, presumably to fuel the increased growth demands during restoration of the damaged tissue.

### The Crypt, Crypt-Villus Junction, and Lower Villus exhibit elevated Signaling during Regeneration

We explored whether the spatial transcriptomics could be used to detect dynamic interactions between cells in distinct spatial domains. Graph-based unsupervised clustering divided the stromal compartments into three domains: the sub-crypt, inter-crypt, and villus stroma (Fig. 4A). Within the control tissues, DGE and subsequent GO term analysis suggested distinct roles for each stromal region (Supplemental table 10). The villus stroma was enriched for adaptive immune-related functions, including MHC-associated and IgG-related gene ontology terms (Fig. 4B), consistent with its role in antigen presentation and local antibody-mediated immune responses at the mucosal interface. In contrast, the crypt and sub-crypt stromal regions were enriched for structural and supportive components, such as collagen production and extracellular matrix organization, highlighting roles in providing mechanical support and modulating epithelial-stromal signaling (Fig. 4B). Quantification of bins across the time course revealed an expansion of the inter-crypt region post-IR, with nearly twice as much area associated with the inter-crypt stromal transcriptome compared to the non-IR state (Fig. 4C). GO analysis between control and post-IR timepoints demonstrated a shift in the villus stroma transcriptome (Fig. 4D-E, Supplemental table 11,12 and 13). Genes elevated during the homeostatic state were associated with functions such as immunoglobulin complex and retinol metabolism signatures. Post-IR there was an increase in genes associated with an active inflammatory response, extracellular signaling, matrix remodeling, and cell adhesion pathways (Fig. 4E), indicative of immune activation and tissue repair processes. Similarly, the inter-crypt stroma transitioned from an adaptive immune signature at day 0, marked by antigen binding and MHC-related terms, to an activated state enriched for complement activation, chemotaxis, cytokine signaling, and innate immune pathways (Fig. 4F), consistent with a transition toward innate immune activation, inflammatory response, and tissue remodeling following injury.

To more specifically interrogate signaling interactions within and between tissue compartments, we utilized CellChat^22^, an algorithm that leverages annotated datasets of receptor-ligand pairs to predict signaling interactions between cells. We used the spatial information in the data to limit potential interactions detected by CellChat to bins within 200µm of each other. A summary of signaling interaction strength for each region at Day 0 and Day 3 revealed a general increase in signaling activity during regeneration, with the crypt and junctional epithelium and the crypt and villus stromal compartments being particularly more active during day 3 (Fig. 4G). By comparison, signaling activity at the villus tip and villus epithelium were similar between the IR and non-IR samples, a trend observed across the independent experiments (Fig. 4G and Supplemental Fig. 4A). Turning a focus to specific cluster-cluster interactions, it was apparent that the strongest gain in interactions at day 3 were occurring between the crypt stroma and crypt epithelium clusters (Fig. 4H). Of all the predicted active signaling pathways that are dynamic between day 0 and day 3 (Fig. 4I and Supplemental Fig. 4C), we decided to investigate those that have documented roles in regeneration after tissue damage, including Transforming growth factor beta (TGFB)^26, 28^, Macrophage Migration Inhibitory Factor (MIF)^38^, neuregulin-1 (NRG)^39, 40^, the complement system^41, 42^, guanylate cyclase C (GUCA)^43, 44^, EGF^45–47^, vascular endothelial growth factor (VEGF)^48^, Angiopoietin-like proteins (ANGPTL)^49^, insulin-like growth factor (IGF)^50^, tumor necrosis factor (TNF)^51^, galectin^52^, collagen^13, 53^, tenascin^54^, prostaglandin E2^55, 56^, and retinoic acid metabolism^57^ (Fig. 4J). We had previously observed a role for TGFB signaling during regeneration^28^ and RNAscope pinpointed *Tgfbr2* and *Tgfb1* expression: *Tgfbr2* was found to be elevated in the crypt and lower villus epithelium and throughout the stroma in day 3, whereas *Tgfb1* was elevated in the stroma at day 3^28^. The spatial analysis confirmed our previous studies, showing similar changes in both *Tgfb* receptor and ligand expression (Fig. 4K-L and Supplemental Fig. 4D). CellChat also predicted signaling from TGFB1 produced in the villus and crypt stroma to be received in the crypt and junction epithelium (Fig. 4I and J Supplemental Fig. 4B). Other signaling pathways upregulated during regeneration and their predicted source and target regions are summarized, such as MIF predominantly produced in the crypt epithelium signaling to the surrounding stroma (Fig. 4M). The results were reproducible across different replicates (Supplemental Fig. 4E-F), giving us confidence to explore more novel results of the CellChat analysis and highlight the utility of this spatial transcriptomics data as a resource for investigating the dynamic expression of genes across the time course of the intestinal damage-repair cycle. The easily accessible dataset should help support research across the field of intestinal regeneration (<u>GSE303705</u>, Supplemental ‘spatial data guide’).

## DISCUSSION

Single-cell transcriptomics has transformed our ability to classify cell types and states, including when studying intestinal regeneration. However, scRNA-seq lacks spatial context, limiting insights into tissue organization and cell-cell interactions. The process of harvesting single cells may also compromise their biology. Traditional in situ hybridization techniques are required to validate single-cell findings and obtain spatial information, however, they have low throughput. The datasets generated in this study overcome these challenges by providing spatially resolved mRNA expression data for over 19,000 genes per experiment. They replicate prior RNAscope results^28^ (Fig. 4L), and recapitulate known biology, including zonation of villus genes at homeostasis^25^ and upregulation of regenerative markers during repair^12, 58^. We also confirm that signaling pathways previously implicated in regeneration exhibit expression of key ligands and receptors in relevant tissues^50, 51^, further validating the approach as an alternative for RNA FISH.

Our dataset can help validate future studies and serve as a resource for new hypothesis testing. For example, we highlight distinct patterns in interferon-signaling pathway target genes (*Isg15*, Fig. 1I) and the capability for intra-tissue differential expression analysis (Fig. 1J). While we confirm known regeneration-associated pathways (Fig. 4I-J), we also identify several novel candidates using the spatial component of CellChat (Supplemental Table 14) that merit further investigation. These spatial transcriptomic datasets offer a time- and cost-efficient strategy to prioritize targets for future studies focused on regeneration, inflammation, and therapeutic discovery.

Spatial assays are rapidly gaining popularity due to their ability to preserve tissue context while enabling deep molecular insights^59–61^. Spatial proteomics allows mapping of protein expression, with resolutions ranging from single-protein detection to ∼20-50µm resolution in mass-spectrometry based approaches^62^ . Spatial metabolomics is also emerging as a powerful tool for mapping metabolite distribution at similar resolution (∼20µm), providing complementary insights into cellular function and disease states^63–65^. Recent advances such as deep visual proteomics integrate imaging, AI, and mass spectrometry to profile disease-relevant cells with high precision^66, 67^. Multiscale approaches that combine targeted and exploratory methods are expanding both the depth and breadth of spatial data^62, 68–71^. Future advances to integrate spatial -omics layers in the gut will help uncover disease mechanisms, inform therapeutic discovery, and advance precision medicine^60, 72^.

Grant Support

The work was supported by grants including the Initiative for Maximizing Student Development (IMSD) Pre-Doctoral Fellow, NIH (T32GM139804), Scholars and Early-Stage Advancement (SEA) Initiative, NIH (3P30CA072720-24S2), NCI R01 and supplement R01CA190558 to MV and RO, the NIH R01DK126446, RC2DK140862, and CA190558 to MPV, Cancer Center Support Grant (CCSG, P30CA072720) from the National Cancer Institute including a New Investigator Award for JY.

Abbreviations

IR (irradiation), DGE (Differential Gene Expression), EdU (5-ethynyl 2’-deoxyuridine), scRNAseq (single-cell RNA-sequencing), MNA (mean normalized average)

## Supporting information

Supplemental table description

spatial data guide

## Acknowledgements

Services, results, and products in support of the research project were generated by the Rutgers Cancer Institute, Immune Monitoring and Flow Cytometry Shared Resource, supported, in part, with funding from the NCI-CCSG P30CA072770-5920, particularly Ankit Saxena, Emmanual Zachariah, Curtis Krier, the Biomedical Informatics Shared Resource, Wenjin Chen, and the Biospecimen Repository and Histopathology Service Shared Resource, Kelly Watkins, supported, in part, with funding from NCI-CCSG P30CA072720-6852. Additionally, by the School of Arts and Sciences Human Genetics Institute of New Jersey Imaging Core. Some illustrations were created using Biorender.com.

## Author contributions

Rachel Ofer; Lead; Conceptualization, Data curation, Formal analysis, Funding acquisition, Investigation, Methodology, Project administration, Software, Supervision, Validation, Visualization, Writing -original draft, Writing review&editing Tyler Tran; Lead; Formal analysis, Software, Validation, Writing -original draft; Supporting; Data curation, Visualization, Data curation Ping He; Supporting; Investigation, Validation, Writing review&editing Xia Qui, Emmanuel Zachariah, Curtis Krier, Ankit Saxena; Supporting; methodology Xinyu Hu, Jiekun Yang; Supporting; Software Michael Verzi; Lead; Conceptualization, Funding acquisition, Methodology, Project administration, Resources, Supervision, Writing -original draft, Writing review&editing

## Data transparency statement

VisiumHD Spatial Transcriptomics data from this publication have been deposited to GEO under the following accession number: <u>GSE303705</u>, and repository URL: https://www.ncbi.nlm.nih.gov/geo/query/acc.cgi?acc=GSE303705. A supplementary tutorial document points the reader to the user-friendly download of the Loupe browser files needed to access the datasets (Supplemental document “Spatial_data_guide”). The code supporting the findings of this study are available upon request.

