## Supplementary figures and images for "A Spatial and Temporal Transcriptomic Atlas of Mouse Intestinal Regeneration"

### Supplemental figure 1

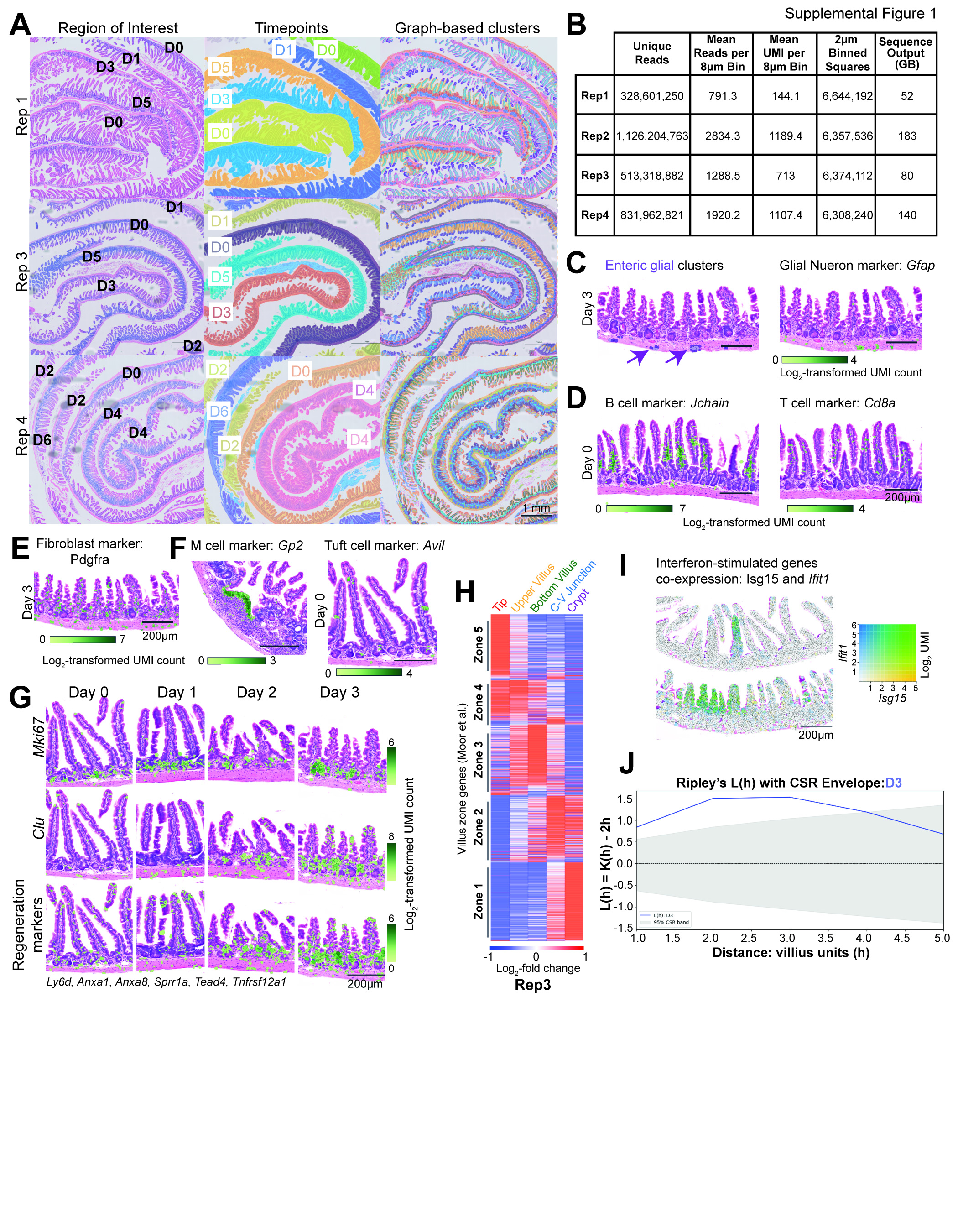

### Supplemental figure 2

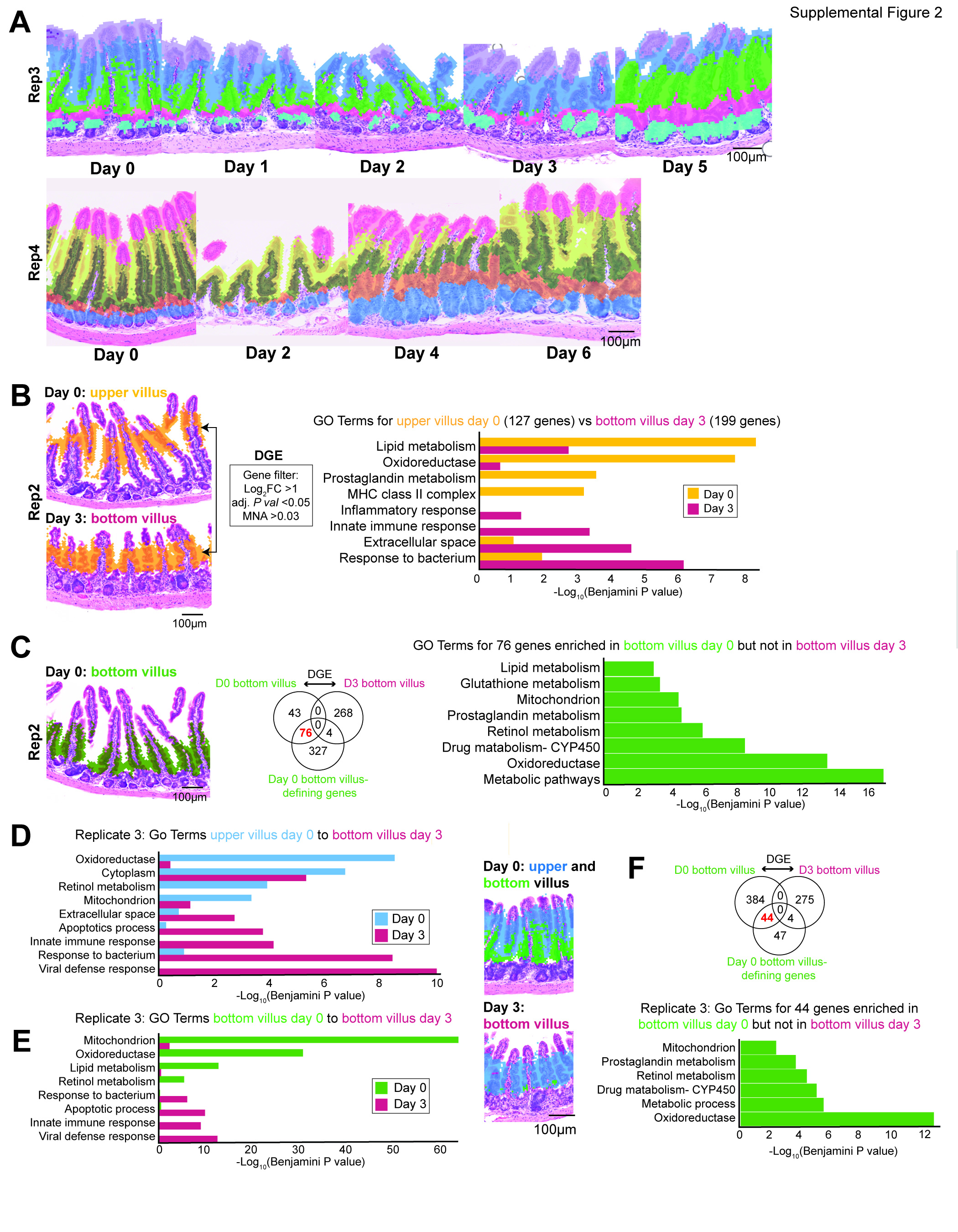

### Supplemental figure 3

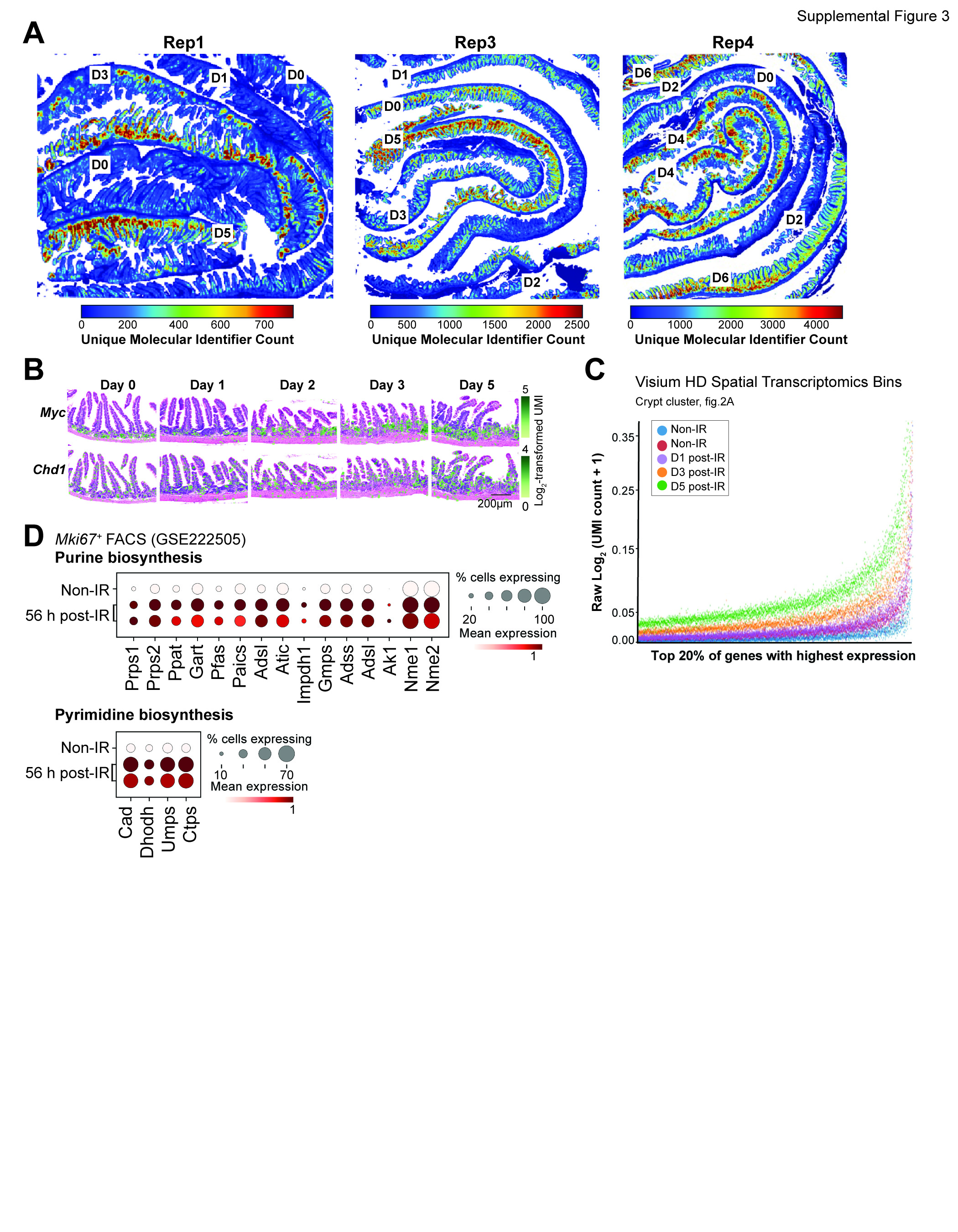

### Supplemental figure 4

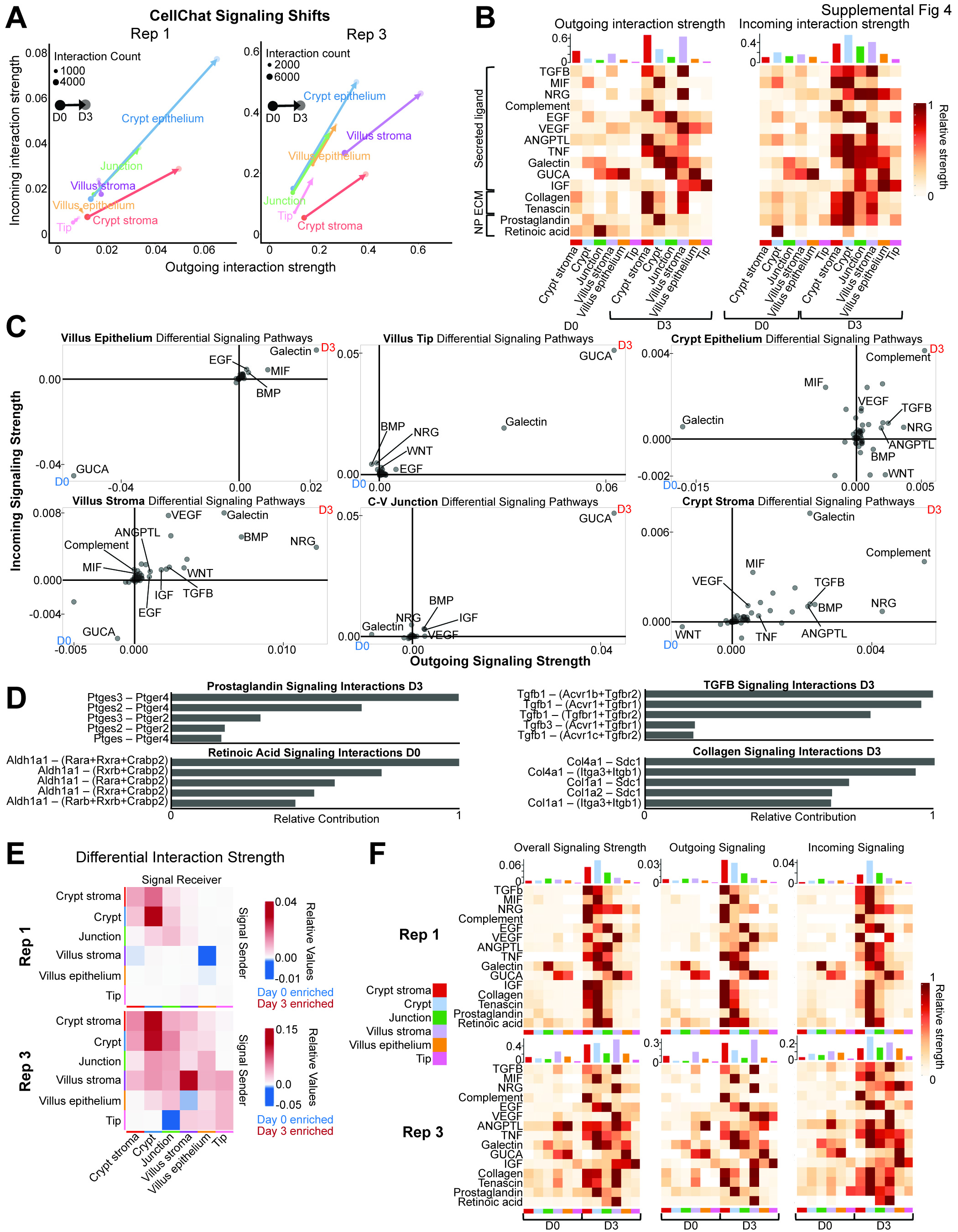
