## Supplemental figure legend for "A Spatial and Temporal Transcriptomic Atlas of Mouse Intestinal Regeneration"

**Supplemental figure 1. Spatial transcriptomics data from multiple replicates confirms consistent clustering and region-specific gene expression.**

**(A)** Spatial capture regions from replicates 1,3 and 4; H&E (left), brushed timepoints (middle) and unsupervised graph-based clustering of 8µm bins (right). Rep1 n = 5; Rep3 n = 6; Rep4 n = 6 independent samples.

**(B)** High-definition spatial transcriptomics data metrics per replicate.

**(C)** Graph-based spatial clusters annotations for enteric glial (left), and enteric glial marker, *Gfap* (right), replicate 2 is shown.

**(D-G)** Spatial-temporal expression ($log_2$-transformed UMI counts) of common markers- B cell (*Jchain*), T cell (*Cd8a*); **(E)** Fibroblasts (*Pdgfra*); **(F)** M cell (*Gp2*); Tuft cells (*Avil*); **(G)** proliferation (*Mki67*), and regeneration markers (*Clu*[12], and sum log2-transformed UMI count for *Ly6d, Anxa1, Anxa8, Sprr1a, Tead4, Tnfrsf12a*).

**(H)** $Log_2FC$ expression of published zone-specific gene sets[25] (1854 genes) across the five epithelial clusters in replicate 3 reflects known anatomical regions along the crypt-villus axis, validating the functional identity of each cluster.

**(I)** The spatial co-expression of *Isg15* and *Ifit1* highlights distinct, region-specific patterns of interferon-stimulated gene activation co-localizing on sparse villi at day 0 time point.

**(J)** The 1D formulation of Ripley's L-function was used to assess the spatial clustering distribution of $Isg15^+$ villi, revealing significant local clustering for day 3 post-IR at distances within one to four surrounding villi. A Monte Carlo simulation assuming random distribution was performed for 1000 random point patterns with the same number of events ($Isg15^+$ villi) as the observed sample, creating a 95% simulation confidence interval for each replicate.

**Supplemental Figure 2. Alterations in villus patterning appear during regeneration across biological replicates.**

**(A)** Unsupervised clustering identifies five transcriptionally distinct epithelial clusters spanning the crypt-villus axis, revealing a shift in zonation patterns that is consistent across replicates: replicate 3 (top) and replicate 4 (bottom).

**(B)** DGE analysis between upper villus cluster of day 0 and bottom villus cluster of day 3 reveals a shift from metabolic functions to immune-related functions, as reflected in representative GO terms.

**(C)** GO enrichment analysis of 76 genes enriched in villus bottom day 0, but not at villus bottom day 3, demonstrating a dramatic loss of villus-bottom associated functions at day 3.

**(D-F)** Analysis of data from replicate 3 confirms the reproducibility of the findings shown in Fig. 2F and Supplemental Fig. 2B-C.

**Supplemental figure 3. Hypertranscription observed in regenerating intestinal tissues across multiple replicates.**

**(A)** Raw UMI heatmaps from spatial replicates 1, 3, and 4 demonstrate elevated transcript abundance post-IR. Each heatmap displays total raw linear UMI counts per barcode, showing global transcriptional upregulation at regenerative timepoints.

**(B)** Expression of two markers of hypertranscription: the universally amplifying transactivator, *Myc,* and chromatin remodeler *Chd1.* Both markers are increased in regenerating tissues post-IR.

**(C)** Global amplification of the transcriptome observed in regenerating tissues. Mean $\log_2$-transformed raw UMI counts of the top 20% most highly expressed genes are plotted across timepoints in replicate 1 spatial data. Data is limited to bins assigned to the crypt cluster based on unsupervised graph-based spatial clustering.

**(D)** Nucleotide metabolism activity is elevated starting at day 3 post-IR. Normalized mean expression dot plots show increased *de novo* purine and pyrimidine biosynthesis gene expression in FACS-sorted *Ki67⁺* epithelial cells at 56 hours post-IR relative to non-IR controls. Data from Chen et al. scRNA-seq dataset (GSE222505)[20].

**Supplemental Figure 4. Replicate analysis confirms regional stromal signaling changes during intestinal regeneration.**

**(A)** Differences in outgoing and incoming signaling strength between day 0 and day 3 post-IR for each stromal region based on CellChat-inferred signaling interactions in replicates 1 and 3.

**(B)** Outgoing and incoming signaling strength by region at day 0 and day 3 post-IR, highlighting pathways previously implicated in epithelial regeneration, including TGFB, complement, and MIF. Results indicate region-specific changes in secretion and reception roles in replicate 2.

**(C)** Differential pathway-level signaling strength between day 0 and day 3 for each stromal domain based on CellChatDB's secreted ligand signaling database. Signaling pathways previously associated with regeneration are annotated.

**(D)** Relative contribution of the top 5 specific genes within the predicated cell-cell signaling pathways of prostaglandin, TGFB, collagen, and retinoic acid signaling, scaled to the strongest interaction per signaling pathway.

**(E)** Comparison of CellChat-inferred interaction networks at day 0 (blue) and day 3 (red) for replicate 1 (top) and replicate 3 (bottom). Consistent regeneration-associated increase in stromal signaling activity was observed.

**(F)** Outgoing, incoming, and total signaling activity by region for select regeneration-relevant pathways across replicates 1 (top) and 3 (bottom).
