## Supplemental table description for "A Spatial and Temporal Transcriptomic Atlas of Mouse Intestinal Regeneration"

**List of Supplemental Tables and Description of Data**

**Spatial Data Guide:**
Guide for downloading Loupe Browser and accessing spatial transcriptomic datasets.

**Supplemental Table 1 (Fig. 1E-G):**
Gene lists for each of the five epithelial zones (villus tip, upper villus, bottom villus, crypt-villus junction, and crypt) at Day 0, identified via differential gene expression (DGE) analysis between zones. Includes gene ontology (GO) terms for each zone. Data shown from replicate 2.

**Supplemental Table 2 (Fig. 1J):**
GO term enrichment analysis comparing *Isg15*$^{+}$ versus *Isg15*$^{-}$ cells on day 0 (Rep2).

**Supplemental Table 3 (Supplemental Fig. 2B):**
DGE and GO term analysis comparing upper villus at Day 0 to bottom villus at Day 3 (Rep2).

**Supplemental Table 4 (Fig. 2F):**
DGE and GO term analysis comparing bottom villus at Day 0 versus Day 3 (Rep2).

**Supplemental Table 5 (Supplemental Fig. 2C):**
DGE and GO term enrichment for genes highly expressed in bottom villus at Day 0 (i.e., genes downregulated by Day 3 that are typically enriched in the bottom villus; Rep2).

**Supplemental Table 6 (Supplemental Fig. 2D):**
GO term analysis comparing upper villus at Day 0 to bottom villus at Day 3 (Rep3).

**Supplemental Table 7 (Supplemental Fig. 2E):**
DGE and GO term analysis comparing bottom villus at Day 0 versus bottom villus Day 3 (Rep3).

**Supplemental Table 8 (Supplemental Fig. 2F):**
DGE and GO term enrichment for genes highly expressed in bottom villus at Day 0 (i.e., genes downregulated by Day 3 that are typically enriched in the bottom villus; Rep3).

**Supplemental Table 9 (Fig. 3I):**
DGE analysis comparing cells with high- versus low-UMI (Unique Molecular Identifier) counts (Rep2).

**Supplemental Table 10 (Fig. 4B):**
DGE and GO term enrichment for stromal subpopulations (villus, inter-villus, and sub-crypt) at Day 0 (Rep2).

**Supplemental Table 11 (Fig. 4E–F):**
DGE and GO term analysis comparing stromal populations (villus) across Day 0, Day 3, and Day 5 (Rep2).

**Supplemental Table 12 (Fig. 4E–F):**
DGE and GO term analysis comparing stromal populations (inter-crypt) across Day 0, Day 3, and Day 5 (Rep2).

**Supplemental Table 13 (Fig. 4E–F):**
DGE and GO term analysis comparing stromal populations (sub-crypt) across Day 0, Day 3, and Day 5 (Rep2).

**Supplemental Table 14 (Fig. 4M):**
Predicted cell–cell interaction data.
