## Supplementary material for "A Spatial and Temporal Transcriptomic Atlas of Mouse Intestinal Regeneration": spatial data guide

Using spatial transcriptomics datasets- a quick guide on getting started

Rachel Ofer, Micheal Verzi


### Introduction

- 10x Genomics Loupe Browser is a visualization tool developed by 10x Genomics and is the default method for visualizing and analyzing the data. LoupeR package can also be used. Both are available here.
- .cloupe file output contains the gene expression matrices, spatial barcode information and image overlays. This file is designed to be opened using Loupe Browser. A .cloupe file is available on GEO for each dataset: GSE303705.

### Installation and setup

To view/access the spatial transcriptomics datasets, you will need to:

1. Download and install the latest version of Loupe browser from 10x Genomics download center
2. Download the dataset from GEO database. These are the .cloupe files
3. Open the .cloupe file with Loupe browser- the dataset is ready to explore!

### Available datasets

Below is a list of available datasets, we recommend starting with Rep2_square_008um_cloupe.cloupe or Rep2_segmented_outputs_cloupe.cloupe

| Cloupe file name | Number of 2µm bins | Mean UMIs per 8µm bin | Genes detected | Timepoints Included |
|---|---|---|---|---|
| Rep1_square_008um_cloupe.cloupe | 6,644,192 | 144.1 | 19,035 | D0, D1, D3, D5, D0 |
| Rep2_square_008um_cloupe.cloupe | 6,357,536 | 1189.4 | 19,037 | D4, D2, D1, D5, D3, D0 |
| Rep3_square_008um_cloupe.cloupe | 6,374,112 | 713 | 19,029 | D4, D2, D1, D0, D5, D3 |
| Rep4_square_008um_cloupe.cloupe | 6,308,240 | 1107.4 | 19,042 | D2, D6, D2 D0, D4, D4 |

Cell segmentation output for rep2 is also available:

| Cloupe file name | Number of cells | Mean reads per cell | Median genes per cell | Median UMIs per cell |
|---|---|---|---|---|
| Rep2_segmented_outputs_cloupe.cloupe | 182,983 | 4,527.3 | 999 | 1502 |

Example of 8x8um vs Cell Segmentation:

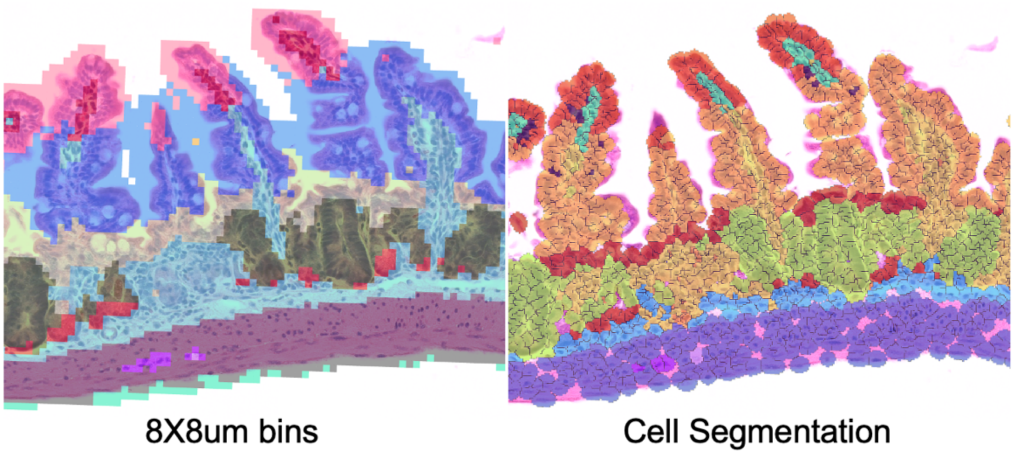


Further resources

Analysis tutorials and comprehensive walkthrough of most features and analysis tools available on loupe browser can be found at the 10x website, here, including analysis using unsupervised clustering, mapping “features” (genes of interest) and cell types, filtering barcodes, running DGE, and using the co-expression feature.

Treatment description per dataset with figures per region of interest (ROI)

1. Rep1 dataset description from outer in inner tissue: Day 0, Day 1, Day 3, Day 5, and Day 0 post-IR.

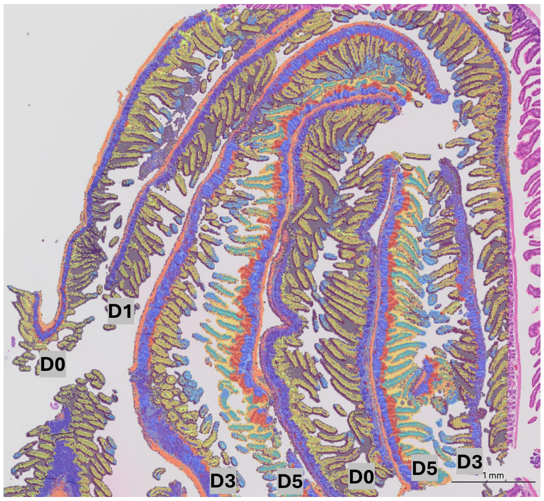

2. Rep2 dataset description from outer in inner tissue: Day 4, Day 2, Day 1, Day 5, Day 3 and Day 0 post-IR.

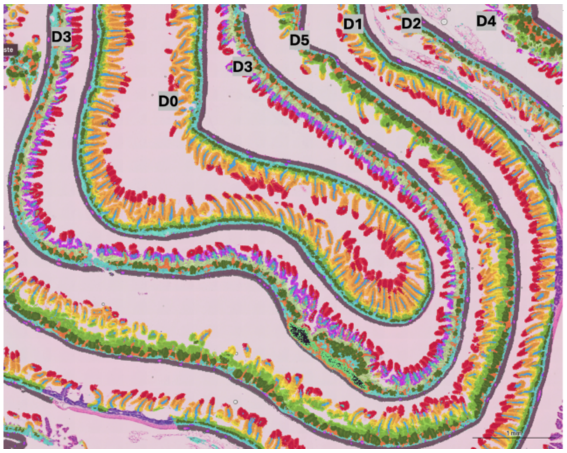


3. Rep3 dataset description from outer in inner tissue: Day 4, Day 2, Day 1, Day 0, Day 5 and Day 3 post-IR.

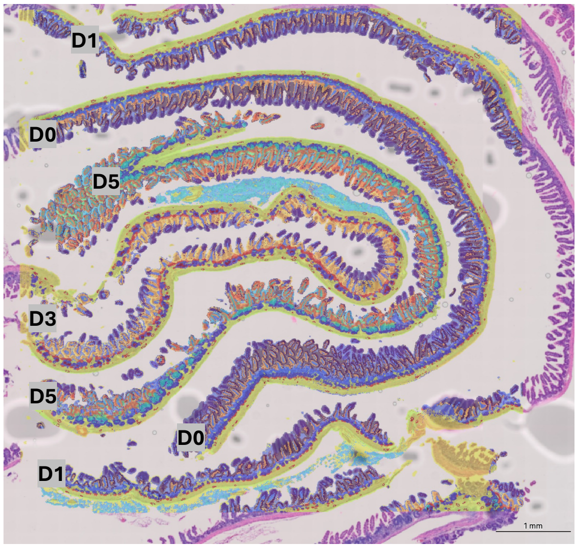

4. Rep4 dataset description from outer in inner tissue: Day 2, Day 6, Day 2, Day 0, Day 4 and Day 4 post-IR.

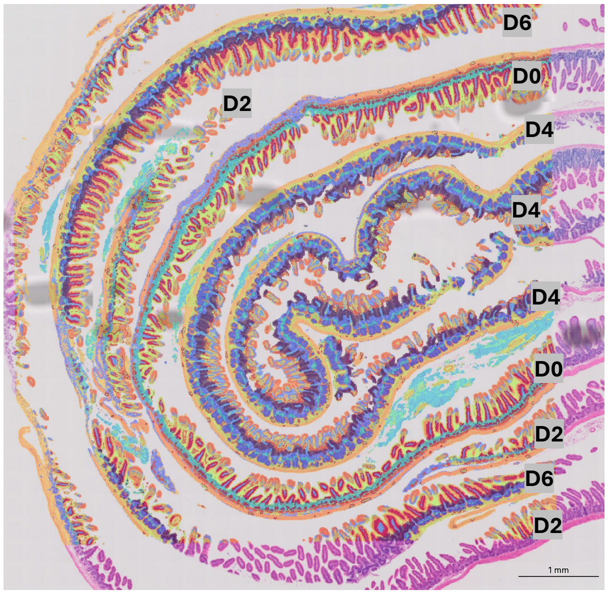
